# *CBX3* Deletion Overcomes Ferroptosis Resistance in Colorectal Cancer by Targeting IFNγ/SLC7A11/SLC3A2 Axis

**DOI:** 10.64898/2026.10.23.761323

**Authors:** Yao Xiang, Yuanji Fu, Christophe Desterke, Eric Batsché, Abdelali Jalil, Rima Elhage, Clémence Duruel, Jean-Christophe Beche, Eric Frapy, David Skurnik, Caroline Reisacher, Sedlik Christine, Eliane Piaggio, Laurence Arbibe, Ahmed Hamaï, Yunhua Chang

## Abstract

System Xc⁻ (SLC7A11/SLC3A2) maintains redox balance by supporting glutathione synthesis, thereby preventing lipid peroxidation and ferroptosis, and represents a promising therapeutic target in cancer. In this work, we revealed that an important epigenetic regulator, CBX3 deletion significantly increased intracellular ROS and lipid ROS level following ferroptosis inducing, particularly under IFNγ stimulation, thereby sensitized CRC cells to ferroptosis *in vitro* using human CRC cells as well as *in vivo* using MC38 syngenetic tumors. RNAseq with mice syngeneic tumors revealed an 11-gene oxidative stress signature downregulated in *Cbx3*KO tumors after combined treatment; its low expression score correlated with improved overall survival in CRC patients. Among them, *Slc7a11*, *Ptgs2*, and *Cth* were identified as CBX3 direct target genes. Additionally, several amino acid transporters functionally coupled to Xc^-^ antiporter, as *Slc3a2*, *Slc7a5, Slc1a5* and *Slc6a9*, all significantly decreased in *Cbx3*KO tumor under RSL3+IFNγ treatment. Mechanistically, CBX3 binds directly to the promoter of *Slc7a11* or *Slc3a2* to sustain their expression and redox balance to resist ferroptosis in CRC. Knockout of *CBX3* in CRC cells significantly impaired RNA polymerase II recruitment to the promoter of *SLC7A11* or *SLC3A2* and reduced SLC7A11 and SLC3A2 expression, an effect that was further exacerbated under IFNγ stimulation but restored by STAT1 deletion. Overall, CBX3 protects CRC from oxidative stress by maintaining the functional integrity of IFNγ/STAT1/SLC7A11/SLC3A2 axis, positioning CBX3 as a potential therapeutic target for enhancing ferroptosis therapies in CRC.

## Introduction

As one of the most prevalent cancers worldwide, colorectal carcinoma (CRC) continues to exhibit limited and often unsatisfactory treatment efficacy despite different treatment strategies continue to be tried ^1^ ^2^ ^3^ ^4^. Chemoresistance remains a major obstacle, undermining the success of conventional therapeutic approaches ^5^ ^6^ ^7^. Ferroptosis—an iron-dependent, regulated form of cell death driven by lipid peroxidation—started to emerge as a promising strategy to overcome apoptosis resistance in CRC ^8^ ^9^ ^10^. As a unique cell death pattern, ferroptosis is mainly caused by the oxidative stress damage and uncontrolled lipid peroxidation resulting from disruption of the system Xc⁻ (SLC7A11)–glutathione (GSH) –GPX4 axis. Among the key tools that enabled the discovery of ferroptosis were the small-molecule compounds Erastin and Ras-selective lethal 3 (RSL3; chemically (1S,3R)-RSL3). The former induces ferroptosis by depleting GSH through inhibition of the system Xc⁻, while the latter directly inactivates glutathione peroxidase 4 (GPX4), thereby inducing ferroptosis.

The cystine–glutamate antiporter system Xc⁻ is composed of the light chain SLC7A11 and the heavy chain SLC3A2. This transporter enables cystine uptake in exchange for glutamate. Once inside the cell, cystine is reduced to cysteine, a precursor for synthesis a major antioxidant, GSH. System Xc⁻ is thus crucial to anti oxidative stress by promoting GSH production, which protects cells from oxidative damage and ferroptosis. SLC7A11 is often upregulated in cancer, enabling tumor cells resist ferroptosis. Targeting SLC7A11 could be a promising therapeutic strategy to enhance ferroptosis or oxidative stress inducing treatment in oncology ^9^ ^11^ ^12^.

It has been reported that Interferon gamma (IFNγ) downregulates the expression of SLC3A2 and SLC7A11 in tumor cells via increased STAT1 binding around the transcription-start region, which impair the uptake of cystine and as a consequence, promoting tumor cell lipid peroxidation and ultimately ferroptosis ^13^. Hence, enhancing and regulating the ability of IFNγ to promote ferroptosis could open new therapeutic perspectives.

CRC is driven not only by genetic mutations (e.g., APC, KRAS, TP53) but also by profound and reversible epigenetic alterations ^14^. Epigenetic therapy has therefore emerged as a promising therapeutic strategy in CRC. Clinically approved DNA methyltransferase inhibitors such as Decitabine and Azacitidine can reverse aberrant DNA methylation and restore tumor suppressor gene expression. In parallel, histone deacetylase (HDAC) inhibitors, including Vorinostat and Panobinostat, are being investigated—mainly in preclinical and early clinical settings—for their ability to remodel chromatin structure and reprogram gene expression. In addition, naturally derived compounds such as curcumin, epigallocatechin gallate, Vitexin and sulforaphane have shown epigenetic modulatory activity in CRC models even precise effects on chromatin-level regulation remain insufficiently characterized ^15^ ^16^ ^17^.

As a key chromatin organizer, chromobox homolog 3 (CBX3, also known as Heterochromatin Protein 1 gamma, HP1γ), is a member of the HP1 protein family in mouse and human. CBX3 is a canonical reader of repressive histone marks, contributing directly to gene silencing when tethered to H3K9-methylated chromatin ^18^ ^19^. However, CBX3 is not restricted to silent chromatin but also actively participates in productive transcription, particularly at gene bodies via physically associated with elongating RNA Pol II and supporting transcription elongation ^20^ ^21^. Moreover, CBX3 co-transcriptionally recruits splicing factors to its target genes, thereby promoting RNA processing as well as alternative splicing ^22^ ^23^. Therefore, as a multifunctional chromatin–RNA interface protein, CBX3 functions in both gene silencing and gene activation, highly depends on chromatin state, binding partners, and genomic location etc…

In our previous work, we revealed that knocking out *CBX3* could increase the CRC sensitivity to IFNγ signaling via directly increased STAT1 transcription ^24^. Furthermore, our integrative analysis has identified several key ferroptosis regulators, particularly system Xc⁻ (including SLC7A11 and SLC3A2) could be direct targets of CBX3 in CRC. All these findings led us to presume that deletion CBX3 would promote tumor cells ferroptosis via affecting lipid peroxidation and increasing IFNγ sensitivity. This hypothesis is confirmed by this work which revealed that CBX3 deletion directly represses *SLC7A11* and *SLC3A2* transcription in a STAT1-dependent manner. This repression is further reinforced under ferroptosis inducer plus IFNγ treatment, and finally increases lipid peroxidation, a functional marker for ferroptosis *in vitro* in human CRC cells as well as *in vivo* in a syngeneic mouse tumor.

Together, our findings highlight that CBX3 plays a critical role in the finetuning regulation in the crosstalk between INFγ signaling pathway and ferroptosis susceptibility. CBX3 is a promising target to improve ferroptosis inducing therapy efficacy in colorectal cancer.

## Materials and methods

### Cell culture and *Cbx3* KO cell lines construction

Human SW480 and HT29 cells were originated from ATCC (CCL-228 and HTB-38). Mouse MC38 cell line was kindly provided by Dr. Eliane Piaggio, as previously described ^24^. The cells were cultured in Dulbecco’s modified Eagle medium (DMEM) supplemented with GlutaMAX (GIBCO, Life Technology) and 10% fetal bovine serum (FBS, Hyclone) with 5% CO2.

The CRISPR/Cas9-mediated *Cbx3*KO cell line, as HT29, SW480 and MC38 cells, were generated as previously descripted^24^ ^25^. The sgRNA used to KO CBX3 in human CRC cell lines is (GAAGAAAATTTAGATTGTCC or GAATATTTCCTGAAGTGGAA). The sgRNA used for KO CBX3 in MC38 is (GAAGAAAATTTAGATTGTCC).

### HT29 *CBX3*/*STAT1* double knockout cell generation

To generate HT29 *CBX3*/*STAT1* double knockout (DKO) cells, the previously established HT29 *CBX3* KO cells were subjected to a lentiviral transduction. Lentiviral particles (from Sigma-Aldrich) encoding a specific sgRNA targeting human STAT1 (sgRNA sequence: 5’-GCAGAGCGCCTGTATTGGG-3’, targeting exon 2) and carrying a puromycin resistance gene alongside Cas9 and tGFP (U6-gRNA:ef1a-puro-2A-Cas9-2A-tGFP,) were used to transduce the *CBX3* KO cells in the presence of 8 µg/mL polybrene for 24h. Selection was performed by fluorescence-activated cell sorting (FACS) based on GFP expression at 48 hours post-transduction. The cells for qPCR analysis were collected 48h after cell sorting (96h after transfection).

### Ferroptosis induction

20,000 HT29 or 30,000 SW480 cells were cultured per well in a 12-well plate. After 72 hours of culture, 200U/ml of human IFNγ were added to the culture medium. Twenty-four hours later, 4 or 8 µM of RSL3 or 5 µM or 10 µM of Erastin were added for an additional 24 or 48 hours, respectively, before analysis. Live cells with different treatments were analyzed by DAPI staining (2.5 μg/mL) to indicate dead cells by flow cytometry.

### Western-blot

The cells or the tumors were washed once with PBS and lysed in 2X Laemmli buffer (100 mM Tris, pH 6.8, 20% glycerol, 4% SDS, 0.05% bromophenol blue, and 10 mM DTT). The lysates were then boiled 5 minutes and subjected to sodium dodecyl sulfate polyacrylamide gel electrophoresis (SDS-PAGE) gel. The tumors samples were sonicated before boiling. After transfer, nitrocellulose membranes were blotted with the following antibodies: anti-HP1γ (IG-2MOD-IG6-AS, Euromedex, 1:2000), anti-β-ACTIN (A3854, Sigma,1:25,000 or 1:50,000), anti-xct-SLC7A11 (12691, cell signaling, 1:1000), anti-SLC3A2 (CAB5702, AssayGenie, 1:1000), anti-COX2 (12282, Cell Signaling, 1:1000), anti-GPX4 (ab125066, Abcam, 1:1000). Primary antibodies were detected with appropriate secondary antibodies conjugated with horseradish peroxidase and the membranes were developed with an enhanced chemiluminescence (ECL) system (ECL detection kit, Thermofisher, MA, USA). The results were quantified by Image J, ACTIN was used as control for protein loading.

### qPCR

RT-qPCR for cells was performed as previously described (Zhang et al. 2020). Primers used for samples are listed in **Table S1**. 36B4 is used as a reference for target gene quantification with the “delta delta Ct” (ΔΔCt) method.

### Immunohistochemistry

Slides were processed on the automaton Leica Bond RX as previously described. Hematoxylin (blue) counterstaining allows the visualization of cell nuclei. The antibody Anti-4 Hydroxynonenal (4-HNE) (ab46544, ABCAM.) was used for MC38 tumors.

### ChIP assay

ChIP experiments were performed as previously described ^22^. In short, HT29 cells were cross-linked with 1% formaldehyde (28908, Thermo scientific) for 10 minutes at room temperature, followed by quenching of the reaction with glycine (26-128-6405-C, Euromedex) at room temperature. After ultrasonic lysis, the cell lysates were incubated overnight at 4 °C with 3 μg of the indicated antibodies: HP1γ (CBX3) (IG-2MOD-IG6-AS, Euromedex, 1:2000), Pol II (pS2:Ab5095, Abcam, 1:500 and pS5: ab5408, Abcam, 1:500), or mouse nonimmune IgG (I-5381, Sigma) as negative control. Beads were washed extensively, and bound material was eluted with 10 mM Tris-HCl, pH 8, 1 mM EDTA, 1% (w/v) SDS at 65 °C for 15 min. Immunoprecipitated DNA was quantified by PCR with primer (see in **Table S1**) sets specific for the indicated CBX3 binding domain. PCR was carried out with an initial denaturation at 95 °C for 30 s, followed by 40 cycles of 95 °C for 5 s and 60 °C for 30 s. Melt-curve analysis was performed from 65 °C to 95 °C at 0.1 °C/s.). Reference background levels were estimated by ChIP with nonimmune IgG. The relative quantification (enrichment) was calculated with the ratio between ChIP with CBX3 and with nonimmune IgG. IgG samples that failed to amplify were treated as below detection and a conservative Ct equal to the run maximum cycle number (40) was used to calculate lower-bound fold enrichment. All values are means ± SEM from 3 independent experiments.

### ROS and lipid ROS measurement

BODIPY 581/591 C11(D3861, Invitrogen), CellROX Deep Red (C10422, life technologies) or CM-H2DCF-DA (C6827, Invitrogen) were used to measure cellular lipid peroxide level or reactive oxygen species (ROS) in live cells, respectively. Cells were seeded in 12 well-plate (30 × 10^3^cells/well). After 48h, cells were stimulated with or without 200U/ml IFNγ, 24h later treated with 0, 4µM, 8µM RSL3, respectively for another 24h and then cells were incubated with 5μM CellROX or CM-H2DCF-DA 30mins, or 1μM Bodipy 581/591 C11 45mins in 37 °C incubator. Then remove the medium and wash three times with DPBS and analyzed them by LSR Fortessa™ cell analyzer (Becton Dickinson, NJ, USA) or by plate reader (TriStar2S 147 LB942/Berthold).

Lipid peroxidation (MDA) Assay kit (ab1118970) was used for tumor tissues’ lipid peroxidation analysis by following the manufacturer’s instructions.

### MC38 syngeneic mouse tumor models

C57BL/6 male mice (7 weeks) were from Janvier-Labs. Animal studies were approved by the ethical committee of Paris Descartes University (authorization number DAP 23-036). After 1 week of settling down, a total of 0.5×10^6^ MC38 WT and *Cbx3*KO cells were injected subcutaneously into C57BL/6 mice at their fourth inguinal nipple. After 1 week, mice started treatment, respectively, three times per week with DPBS (Control), 5000U of recombinant murine IFNγ (315-05, Pepro Tech), 40 mg/kg RSL3, and 40 mg/kg RSL3 with 5000U of recombinant murine IFNγ, respectively. Ferroptosis inducer (1S,3R)-RSL3 (RSL3, HY-100218A) was purchased from MedChemExpress (MCE, USA). For in vivo experiment, RSL3 was dissolved in buffer contain 10%DMSO 40%PEG400 5% Tween-80 45%Saline. For the mice treated with RSL3 and IFNγ, the first injection of IFNγ is 4 h before the RSL3 injection and the following injections were done at the same time. Tumor size was measured three times per week in two dimensions, the longest tumor dimension (L, length) and the value at right angles to it (W, width), with a caliper. Tumor volume was calculated as an ellipse (tumor volume = (width2 * length)/2). After 16 days treatment, the mice were sacrificed and the tumors were separated and weight.

### RNA sequencing and bioinformatic pipelines

Total RNA was extracted from subcutaneous MC38 tumor at the end of treatment. Each group (Ctrl, RSL3, and RSL3 plus IFNγ) has 3 samples. The tumor was homogenized in 1 ml trizol by a homogenizer (Fisherbrand™ Bead Mill 24), then RNA was extracted following the manufacturer specifications. Total RNA library preparation and sequencing were performed using the DNBseq system at BGI Genomics Co., Ltd. To obtain clean reads, the raw reads were filtered following sequencing using the SOAP nuke parameters: -n 0.001 -l 20 -q 0.4 --adaMR 0.25 --polyX 50 --minReadLen150. Gene quantification used STAR 2-pass alignment on MM10 genome follow by feature counts quantification for the pipeline (39163 annotated genes). The raw count matrix was normalized with EdgeR TMM method follow by voom transformation and filtration of 13283 well-expressed genes.

### Survival analyses with integration of TCGA transcriptomic

Clinical and transcriptomic data were obtained from The Cancer Genome Atlas (TCGA) for CRC ^26^, including colon adenocarcinoma (COAD) and rectal adenocarcinoma (READ) cohorts were downloaded on Cbioportal website ^27^. RNAseq data quantified by RSEM V2 in transcript per millions (TPM) were used to quantified gene regulation ^28^. Patients’ clinical data was associated to expression of selected genes: *CHAC1, TRIB3, FOSB, SESN2, NR4A3, SLC7A11, CTH, PTGS2, CTGF, ALDH1L2,* and *EDN1* to perform overall survival analysis in R software version 4.4.2 with survival R-package version 3.8-3 ^29^. Loop of univariate cox analysis on each gene expression was done with loopcolcox R-package available at the following web address: https://github.com/cdesterke/loopcolcox (accessed on 2025, September 8th). Expression score was computed by extracting Cox beta coefficient and matching them with expression of each eleven markers. Overall survival threshold determination on expression score, and Kaplan Meier analyses were performed with survminer R-package version 0.50.A multi-variable overall survival Cox model was fitted including the expression score, tumor, primary tumor site, age, and sex. Model performance and proportional hazards assumptions were evaluated using cox.zph function ^30^.

### Statistical analysis

For biological experiments, paired, unpaired, or multiple two-sided t-test and two-way ANOVA were performed with the GraphPad Prism based on at least three independent experiments or mice samples according to different experiment conditions. For syngeneic mouse tumor model, treatment groups were assigned in a randomized fashion. For clinical analysis, sample size of the original TCGA COAD transcriptome cohort was taken in account with exclusion of samples comporting missing data which represent less than 0.3 percent of the total cohort. No randomization was performed during the observational studies. The two-sided t-test was applied for clinical co-expression analysis and the log-rank test of the Cox proportional Hazard model was applied for overall survival analysis. For all analysis, P-value inferior to 0.05 was considered as significant.

## Results

### *Cbx3* deletion sensitizes MC38 CRC tumor to ferroptosis inducer RSL3 under IFNγ stimulation *in vivo*

Accumulating evidence has demonstrated that IFNγ promotes lipid peroxidation and ferroptosis in tumor cells *in vivo*^10^. Our previous studies have also identified CBX3 as a negative regulator of IFNγ signaling, suggesting that the presence of CBX3 may limit tumor cell susceptibility to IFNγ-promoted ferroptosis. Based on these findings, we hypothesized that depletion of CBX3 would increase the susceptibility of colorectal cancer (CRC) tumors to ferroptosis inducer, particularly in the presence of IFNγ. We therefore tested this hypothesis using MC38 tumor models *in vivo* under the treatment of IFNγ or/and RSL3 (RAS-selective lethal 3), a potent ferroptosis inducer which directly binds and inactivates GPX4.

WT or *Cbx3*KO MC38 cells were subcutaneously implanted into the flanks of C57BL/6 mice. The mice were divided as control group (Ctrl group), RSL3 group, IFNγ group and Combo group (RSL3 combined with IFNγ) and treated from 7 days after tumor cells implantation.

In MC38 WT tumors, no significant difference was identified in tumor growth and final weight among the four treatment groups (**Figure 1A, left and Figure 1C**). Similarly, in the MC38 *Cbx3*KO group, RSL3 or IFNγ treatment alone did not result in a significant decrease of tumor size compared to the control group (**Figure 1A, middle**). However, combined RSL3 and IFNγ treatment (combo group) resulted in a significant decrease in MC38 *Cbx3*KO tumors volume, as well as final tumor weight compared to other KO groups (Ctrl, RSL3 and IFNγ) or WT combo group (**Figure 1A, middle and right; Figure 1B-1C**). These findings demonstrated that *Cbx3* deletion makes CRC more susceptible to ferroptosis therapy under IFNγ stimulation.

**Figure 1:**
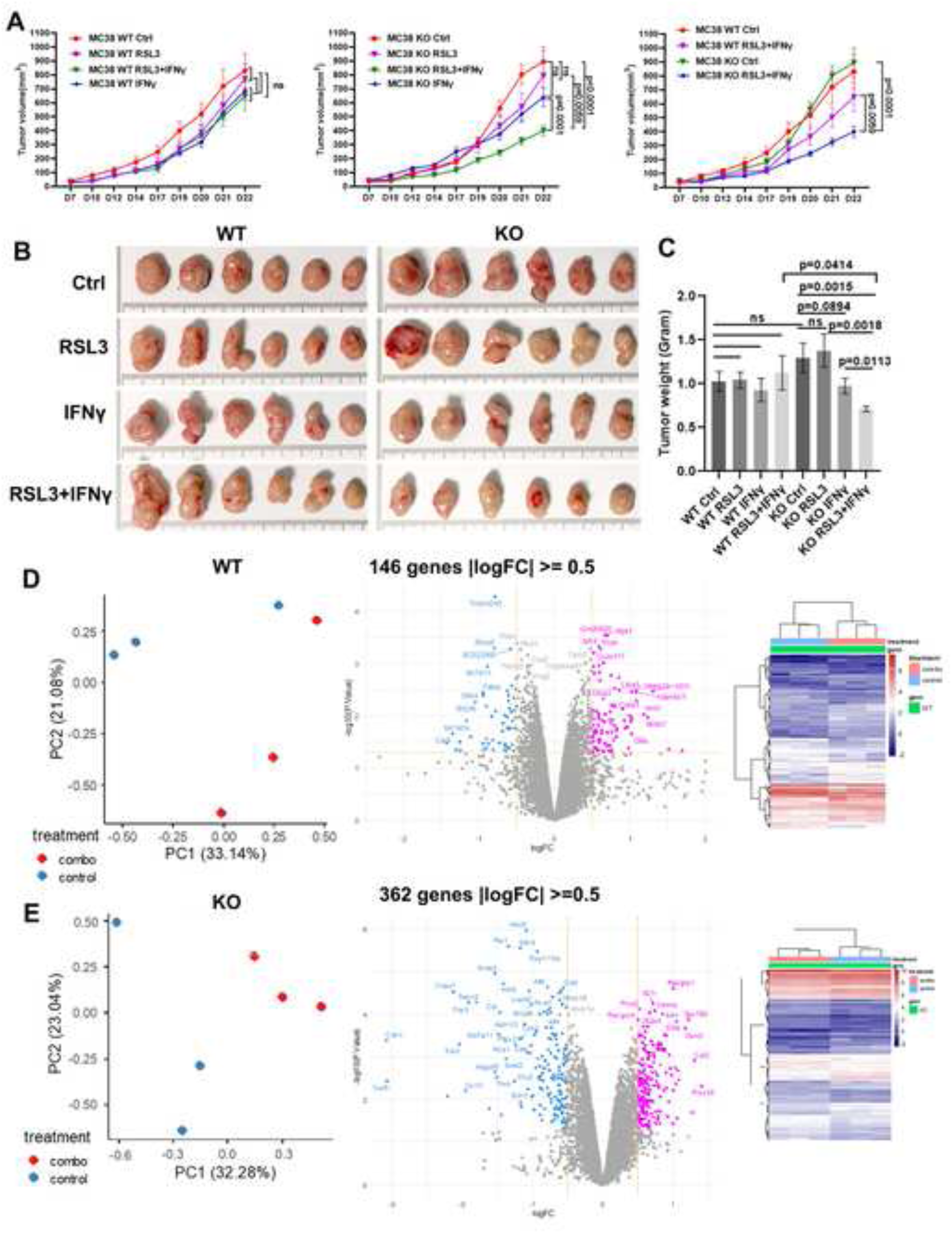
**A**) Left: WT tumors do not present a significant difference between different treatment group (as RSL3, IFNγ or RSL3 + IFNγ) and Ctrl group. **Middle:** for MC38*Cbx3* KO tumors, RSL3 + IFNγ (combo) treatment significantly reduced their size compared to IFNγ, RSL3 and Ctrl group. No significant difference is detected between RSL3 and Ctrl group or between IFNγ and Ctrl group. **Right:** the size of *Cbx3*KO tumors under RSL3 + IFNγ treatment is significantly decreased compared to WT tumors under RSL3 + IFNγ treatment. Two-way ANOVA was applied for statistical analysis, n = 7-8 mice. The error bar represented SEM. **B)** Representative images of the tumors from different treatment groups at D22. **C)** Tumor weights at D22 of different treatment groups. The *Cbx3*KO tumors under combo treatment are significantly lighter than WT combo group (n = 9-13, two-sided t test). The data are presented as means ± SEM. **D and E)** RNA-sequencing (RNAseq) analysis on WT (**D**) and *Cbx3*KO (**E**) MC38 tumor tissues. **Left**): PCA (principal component analysis) visualized gene expression similarity between samples (Combo and Ctrl group in WT and *Cbx3KO* tumors, each group 3 samples) by projecting the data onto the first principal components (PC1 and PC2). **Middle and Right):** Two Volcano Plots combining fold change and p-value to visualize significantly differentially expressed genes in combo group compared to Ctrl group in WT and *Cbx3*KO MC38 tumor. 362 DEGs (differentially expressed genes) with absolute value of Log2FC≥0.5 were identified in *Cbx3*KO tumors (combo vs Ctrl) but only 146 DEGs with absolute value of Log2FC≥0.5 were found in WT tumor (combo vs Ctrl).

### IFNγ combined with RSL3 perturbes diverse metabolic pathways feed into cellular redox balance in *Cbx3*KO CRC tumor

To further investigate the molecular mechanisms underlying this enhanced therapeutic response, we carried out a bulk tumor RNA-sequencing (RNAseq) analysis with WT and *Cbx3*KO MC38 tumors at the end of treatment (D22) (**Figure 1D-1E**). Since the therapy efficacy only appeared in the combo group (RSL3 and IFNγ) for *Cbx3*KO tumor, we thus focused on comparing the combo group with the control group in both KO and WT tumors. Interestingly, RNAseq analysis showed that RSL3 and IFNγ combination therapy induced a more evident gene expression modification in KO than in WT tumor (**Figure 1D-1E**). Compared to their respective control groups, 362 DEGs (differentially expressed genes) with |Log2FC|≥0.5 were identified in *Cbx3*KO tumors, whereas only 146 DEGs with |Log2FC|≥0.5 were found in WT tumor.

Correspondingly, gene set enrichment analysis (GSEA) based on RNAseq data revealed significant metabolic and redox reprogramming in KO tumors under combined RSL3 and IFNγ therapy, characterized by suppression of pathways involved in fatty acid metabolism, cholesterol homeostasis, glycolysis, heme metabolism, and mTORC1 signaling, alongside decreased hypoxia but enrichment of reactive oxygen species–associated pathways. (**Figure 2A-2B**). However, the WT combo group did not exhibit such a reduction, which only exhibited a significant attenuation of oxidative phosphorylation pathways compared to the WT control group (**Figure 2A and Figure S1)**.

**Figure 2.**
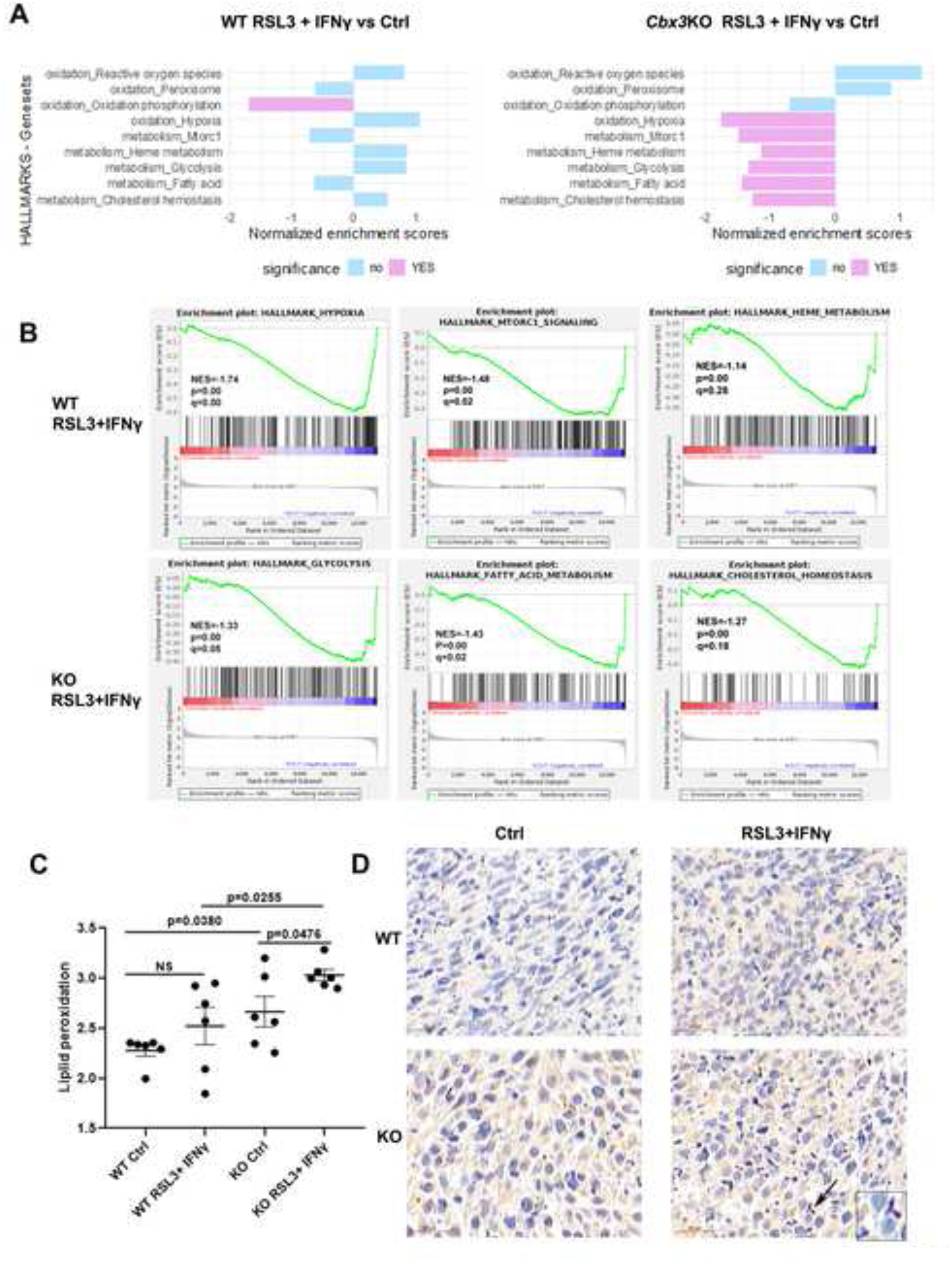
**A**) HallMARK Geneset GSEA analysis based on WT and KO MC38 tumor RNAseq data revealed that combined RSL3 and IFNγ therapy significantly perturbed different metabolic pathways in KO tumors. Signal2Noise uses the difference of means scaled by SD with FDR (false discovery rate) adjustment for multiple testing, n = 3. **B)** For RSL3+ IFNγ group compared to Ctrl group, the 3 most evidently decreased enrichment metabolic pathway in WT or in KO tumors. **C)** The lipid peroxidation level is significantly increased *Cbx3*KO tumors compared to WT tumor. RSL3 and IFNγ treatment further increased the lipid peroxidation level in *Cbx3*KO tumor compared to KO Ctrl tumors. Two-sided Student’s t-test, n=6. The error bar represented SEM. **D)** Immunohistochemistry revealed an increased 4-HNE positive cells in *Cbx3*KO tumor under RSL3+ IFNγ treatment compared to KO or WT ctrl and WT combo group. Hematoxylin (blue) counterstaining allows the visualization of cell nuclei. Scale bar: 20 µm for main image and 5 µm for insert image.

In KO combo group tumors, the negative enrichment of pathways related to fatty acid metabolism, cholesterol homeostasis, glycolysis, heme metabolism, and mTORC1 signaling indicated a coordinated suppression of metabolic and anabolic programs. Concurrently, hypoxia-related gene sets were negatively enriched, suggesting reduced activation of hypoxia-driven adaptive responses. In contrast, gene sets associated with reactive oxygen species were positively enriched, pointing to increased oxidative stress and altered redox regulation. Collectively, these GSEA results highlight a global metabolic reprogramming characterized by diminished energy production and growth signaling alongside enhanced oxidative stress responses between KO combo and Ctrl group. These changes are consistent with decreased tumor growth but increased susceptibility to ferroptosis induction of KO tumor under combo treatment.

### Deletion of *Cbx3* increases lipid peroxidation in CRC tumors, especially under combined IFNγ and RSL3 treatment

We then checked lipid peroxidation level, a very important hallmark of ferroptosis, in CRC tumors with or without IFNγ and RSL3 treatment. The lipid peroxidation level was significantly increased in the KO tumor than in WT tumor. The combo therapy further enhanced lipid peroxidation in KO tumors. However, no significant modification was found between the combo and control group in WT tumor (**Figure 2C**). As a definition marker of ferroptosis, increased lipid peroxidation level suggested an increased lipid ROS stress and ferroptosis susceptibility in KO tumor, especially under RSL3 and IFNγ combo treatment.

At the same time, we also checked 4-hydroxy-2-nonenal (4-HNE), another major marker of cellular lipid peroxidation and oxidative stress, by immunohistochemistry analysis. Once more, under RSL3 and IFNγ treatment, KO tumors exhibited an increased 4-HNE positive cells marking compared to WT tumors (**Figure 2D**).

### Integrated bioinformatics analysis identifies *Slc7a11, Ptgs2* and *Cth* as potential CBX3 targets among the most downregulated genes in KO tumors under IFNγ and RSL3 treatment

Furthermore, RNAseq analysis identified that RSL3 and IFNγ combination therapy induced 47 top DEG with |Log2FC|≥1 in *Cbx3*KO tumors (combo vs Ctrl) and only 20 appeared for WT combo vs Ctrl tumors (**Figure 3A-3B and Figure S2**). Among the top downregulated DEG with Log2FC≤ −1 in KO tumor (combo vs Ctrl group), we identified a STRING network composed of 11 genes. The 11 genes are not among the 20 top DEG in WT tumors and have no intersection with the shared top DEG between WT and KO (combo vs Ctrl) (**Figure 3B-3C**). Metascape pathway and process enrichment analysis revealed that the most evidently enriched term across these 11 genes is response to oxidative stress (**Figure 3D**). Therefore, this cluster with 11 genes is referred as the OS signature in the following text.

**Figure 3.**
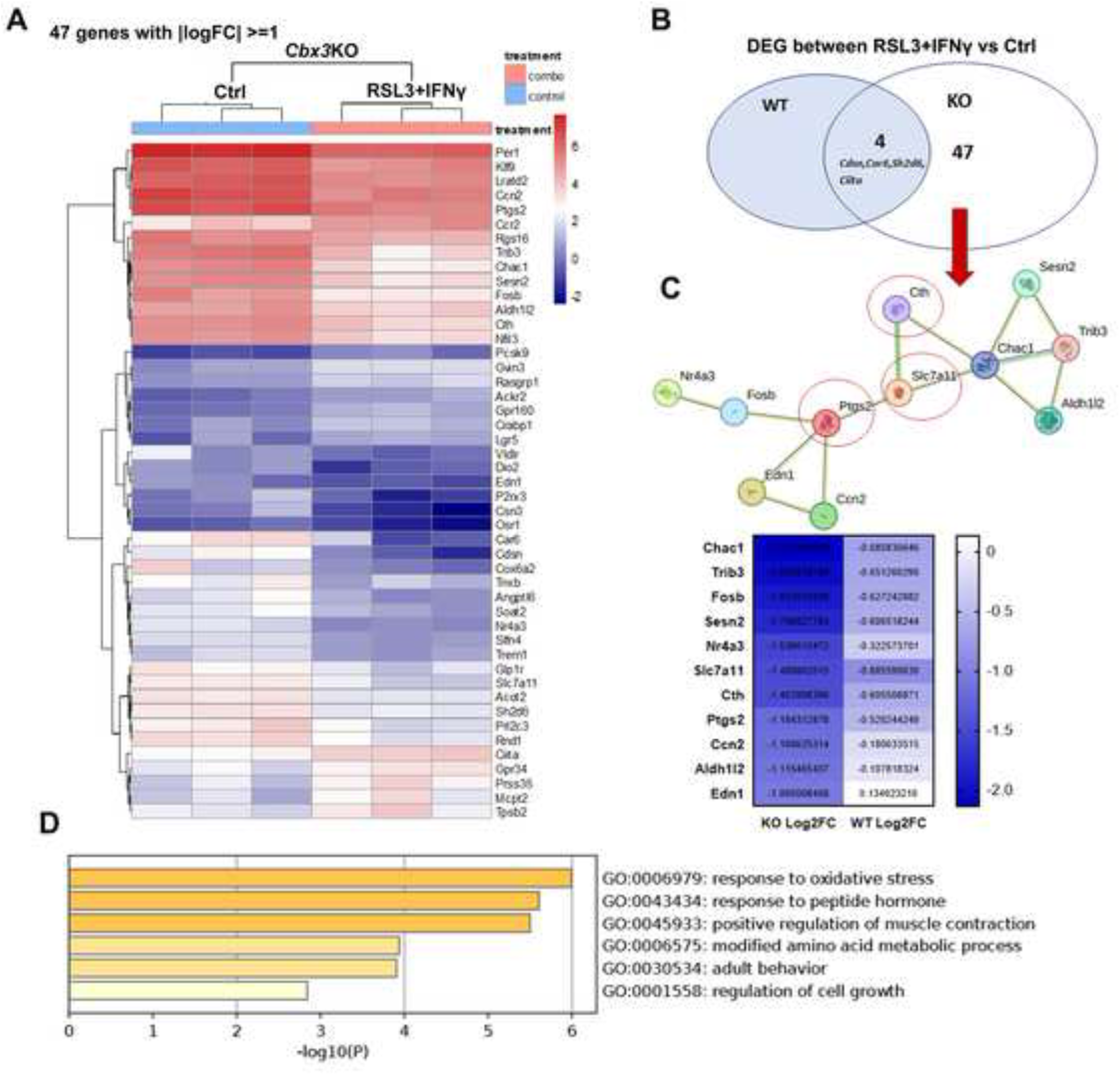
**A**) Heatmap generated from RNAseq data exhibited the differentially expressed genes (47 DEGs) with the absolute value of Log2FC≥1 in KO combo group compared to KO Ctrl group. **B)** Between combo and Ctrl group, only 4 common DEG with lLog2FCl≥1 between WT and KO MC38 tumor: 3 downregulated (*Cdsn*, *Car6* and *Sh2d6*), 1 upregulated (*Ciita*). **C)** Among the most significantly downregulated genes(Log2FC≥-1) between the KO combo group and the KO control group, 11 genes formed a STRING network. **D)** Metascape pathway and process enrichment analysis revealed that 11 genes were enriched in response to oxidative stress and referred as OS signature in the later text. Pathway and process enrichment analysis have been carried out with the following ontology sources: GO Biological Processes, KEGG Pathway, Reactome Gene Sets, and WikiPathways. Terms with a p-value < 0.01, a minimum count of 3, and an enrichment factor > 1.5 (the enrichment factor is the ratio between the observed counts and the counts expected by chance) are collected and grouped into clusters based on their membership similarities.

Furthermore, integrative analysis by crossing the 11 genes in OS signature with the CBX3 target genes identified with a published HCT116 CBX3 ChIP-seq data ^23^, *Slc7a11*, *Ptgs2* and *Cth* were recognized as potential CBX3 target genes as a CBX3 binding peak is discovered in the zone of every gene’s promotor (**Figure 3C** and **Figure S3)**.

### The expression of *Slc7A11*, *Ptgs2* or *Cth* is decreased in *Cbx3*KO CRC tumor, particularly under RSL3 and IFNγ therapy

At the same time, by using Integrative Genomics Viewer (IGV) on the MC38 RNAseq data, an evident decrease in *Slc7a11*, *Ptgs2* and *Cth* transcription levels was observed in KO compared to WT tumors, this diminution becomes even more pronounced after combo treatment (**Figure 4A-4C, see also Figure S4**). The results were then confirmed by western blot analysis, which revealed the protein level of SLC7A11or PTGS2 is significantly reduced in *Cbx3*KO tumors compared to WT tumor, either for Ctrl or for combo groups (**Figure 4D**).

**Figure 4.**
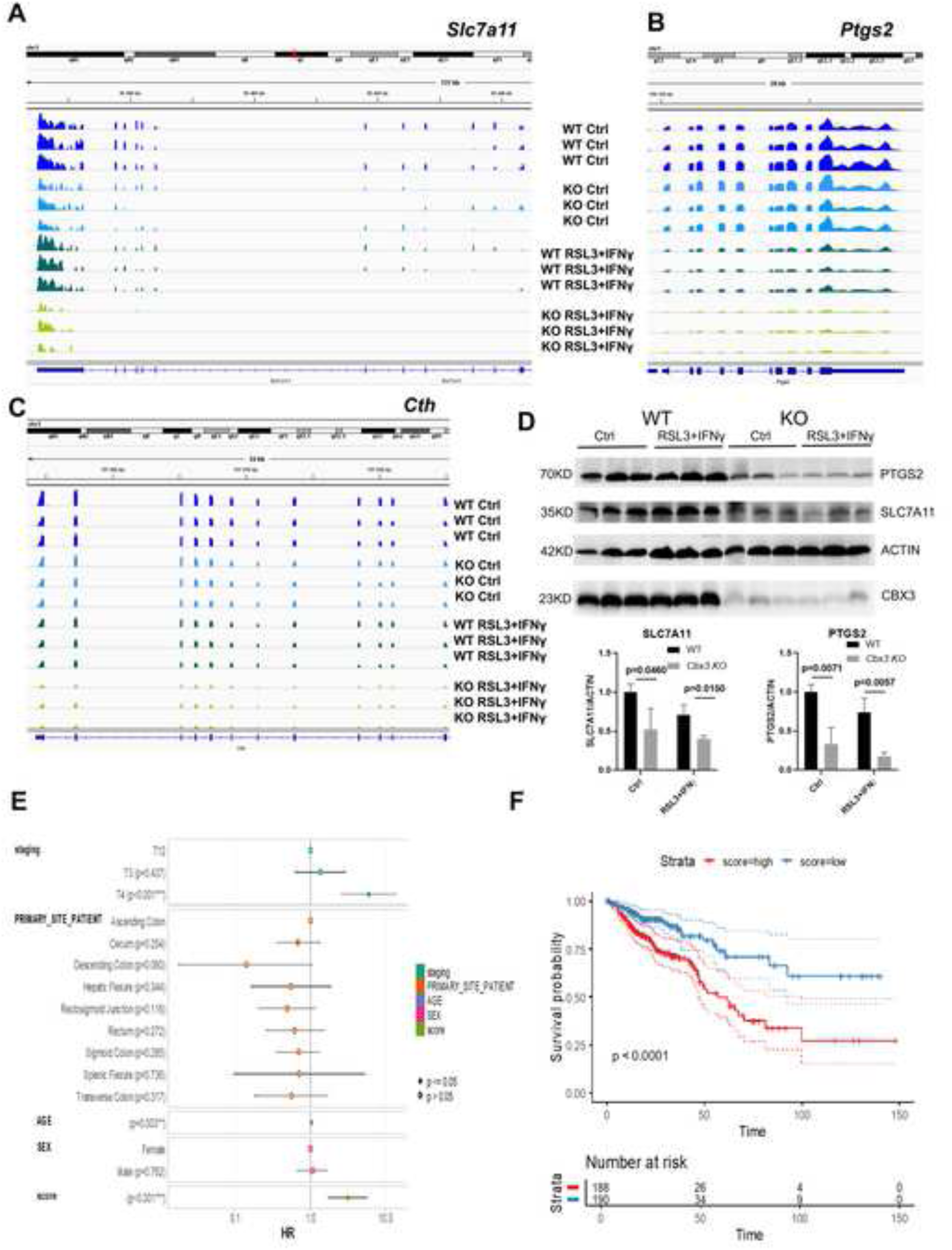
**A, B and C)** Integrative Genomics Viewer (IGV) was applied on MC38 Cbx3KO and WT tumors RNAseq data to visualize Slc7a11, Ptgs2, and Cth transcription levels across the Ctrl group and combined treatment group (RSL3 + IFNγ). **D) Up)** Western blot analysis showed the decreased protein levels of Slc7a11 and Ptgs2 in Cbx3KO MC38 tumor compared to WT tumors in Ctrl and Combo (RSL3+ IFNγ) group. **Down)** Image J was used for the protein level quantification, which revealed a significant decreased of SLC7A11 and PTGS2 expression in Cbx3KO MC38 tumor compared to WT tumors in both Ctrl and Combo (RSL3+ IFNγ) group. Three tumors for each group, two-sided Student’s t-test. The error bar represented SEM. **E)** The OS signature comprising 11 tops downregulated DEG (Log2FC≥-1) between KO ctrl and combo group, whose expression scores demonstrate independent prognostic value in CRC patients beyond standard clinical factors. The forest plot showing hazard ratios (HR) and 95% confidence intervals for the 11 gene score across multiple TCGA cancer types. Significant associations (p < 0.05) are marked with asterisks. Variables such as age, sex, and mismatch repair status (MMR) are included as covariates. **F)** Prognostic impact of the OS signature across survival stratification. Kaplan–Meier survival curves comparing patients with high (red) versus low (blue) expression scores. The score was dichotomized using an optimal cutpoint method. A significant difference in overall survival is observed (log-rank p < 0.0001). The table below indicates the number of patients at risk over time (month) in each group.

SLC7A11 and CTH contribute to GSH production and redox homeostasis, while PTGS2 deregulation is associated with redox imbalance. CBX3 binds to the promoter of *Slc7a11*, *Ptgs2* and *Cth* to maintain their transcription and deletion CBX3 considerably decreased their expression under RSL3 and IFNγ treatment. Combined, these results suggested the potential molecular mechanism underlying the enhanced ferroptosis therapeutic response in KO tumor.

### The low expression score of the 11 genes in the OS signature is linked to better overall survival in CRC patients

To evaluate the prognostic relevance of the OS signature, we performed multivariate Cox regression analyses across CRC transcriptome data from TCGA consortium. A Cox model risk score based on the quantitative analysis of 11 genes was calculated and integrated into a multivariate overall survival model. The forest plot (**Figure 4E**) reveals that the 11 genes expression score is significantly associated with overall survival in different colorectal tumor subtypes. Age, sex, and tumor stages were included as covariates and showed variable contributions depending on the cancer context with linear distribution of their Cox residual (**Figure S5**). Notably, the expression score remained a strong prognostic factor after adjustment for all clinical variables. To further assess the prognostic value of the score, patients were stratified into high and low score groups using an optimal cutpoint method. Kaplan–Meier analysis demonstrated a significant survival difference between the two groups (log-rank p < 0.0001), with the high-score group exhibiting markedly reduced overall survival (**Figure 4F**).

These findings correspond to the increased sensitivity to IFNγ and RSL3 therapies in KO MC38 tumor, support that the signature of these 11 down-regulated genes in KO combo group could be used as a CRC prognostic biomarker and highlight its potential for risk stratification in clinical settings.

### The expression of SLC3A2 and other amino acid transporters functionally coupled to SLC7A11/GSH axis is substantially reduced in *Cbx3*KO tumor under IFNγ plus RSL3 treatment

As the main intracellular antioxidant to protect cells from oxidative stress and maintains redox balance, GSH is synthesized in the cytosol from glutamate, glycine, and cysteine, whose cellular uptake relies on several coordinated amino acid transporters. RNAseq data revealed that IFNγ plus RSL3 treatment reduced significantly the expression of several important amino acid transporters functionally coupled to SLC7A11/GSH regulated redox balance and ferroptosis in *Cbx3*KO MC38 tumors (with Log2FC ≤ −0.5) (**Figure 5A-5C**).

**Figure 5.**
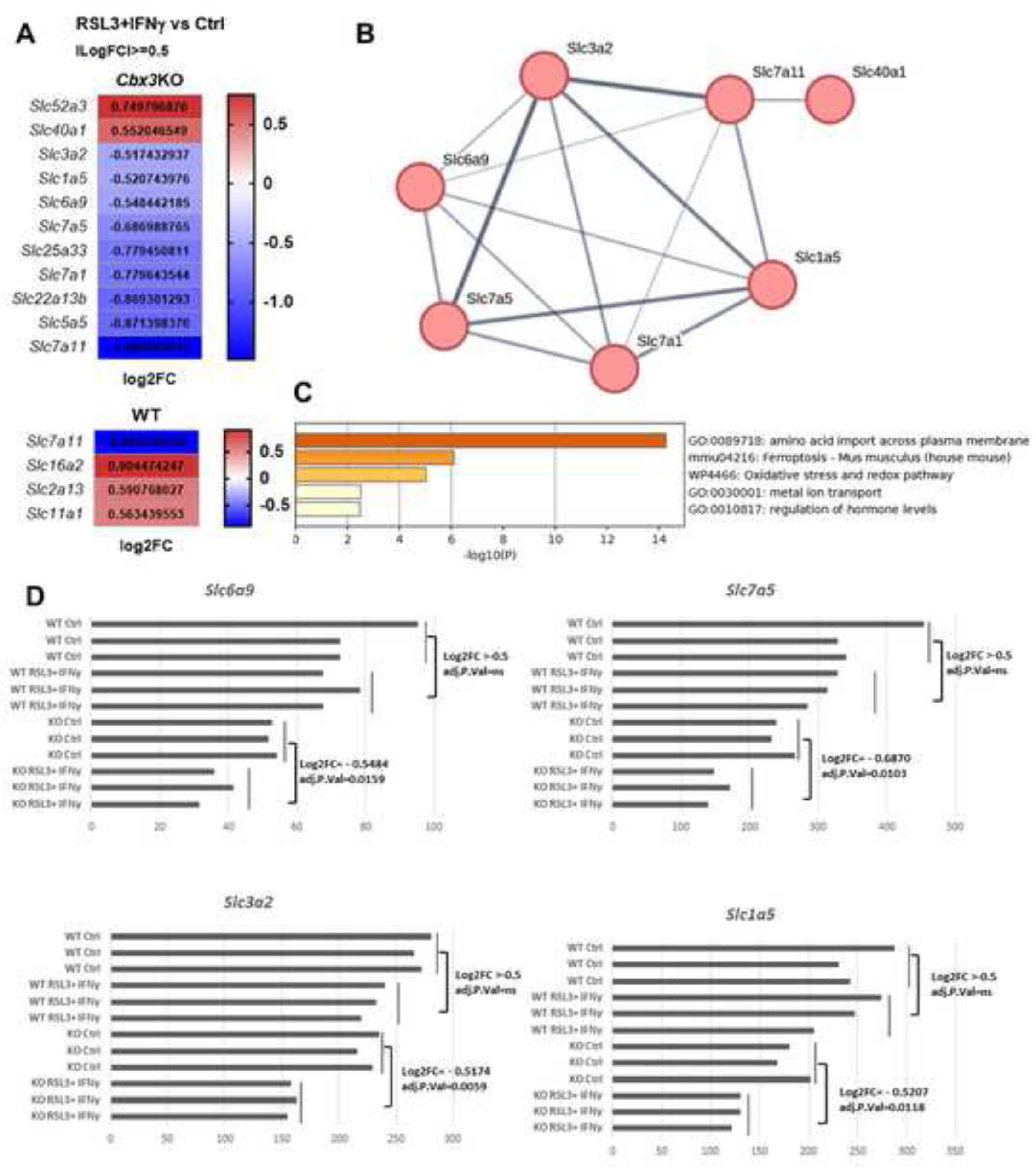
**A**) besides SLC7A11(Log2FC=-1.48), several key amino acid transporters functionally coupled to SLC7A11/glutathione axis are significantly decreased in the KO combo therapy group compared to control group (Log2FC < −0.5, p<0.05). However, these amino acid transporters genes are not identified in DEG with lLog2FCl >=0.5 in WT combo vs WT control group except SLC7A11(Log2FC=-0.88). **B)** A function coupled string network is formed by the identified amino acid transporters. **C)** Metascape pathway and process enrichment analysis revealed that these amino acid transporters enriched in ferroptosis, oxidative stress and redox pathway. **D)** The RNAseq data (normalized by EdgeR) revealed the transcription levels of Slc3a2, Slc1a5, Slc7a5 and Slc6a9 in MC38 Cbx3KO and WT tumors across the Ctrl and the combo group.

The most downregulated amino acid transporter between KO combo and Ctrl group is *Slc7a11* (Log2FC= −1.48), the light-chain of cystine/glutamate transporter system Xc⁻ which primary function is to import extracellular cystine in exchange for glutamate and thereby supports intracellular cysteine availability and glutathione (GSH) synthesis (**Figure 5A**). Moreover, SLC3A2, the essential chaperone for recruiting SLC7A5 and SLC7A11 to the plasma membrane to enable their transport functions, is also significantly downregulated between KO combo and Ctrl group (**Figure 5A-5B, 5D**). Most notably, the HCT116 CBX3 ChIP-seq data ^16^ revealed also a CBX3 binding peak in the promoter of *Slc3a2,* suggesting besides *Slc7a11*, *Slc3a2* could be also a direct target gene of CBX3(**Figure S3**).

Moreover, Alanine-Serine-Cysteine Transporter 2 (ASCT2/SLC1A5) and L-type amino acid transporter 1 (LAT1/SLC7A5), two amino acid transporters mediating the influx of extracellular glutamine required for the optimal function of Xc⁻ system activity, are also significantly decreased in *Cbx3*KO MC38 tumor under combo treatment (**Figure 5D**). Finally, the decreased expression of a major glycine transmembrane transport, SLC6A9 (Glycine transporters 1) suggested an additional aberrant glycine uptake/transport in KO combo therapy group (**Figure 5D**).

In contrary, for WT MC38 tumor, except *slc7a11*(Log2FC=-0.88, combo VS Ctrl group), the others amino acid transporters are not significantly diminished in WT MC38 tumors after combo treatment (**Figure 5A**).

Overall, all the results suggested CBX3 as a key regulator to maintain the expression of different key amino acid transporters, which play crucial role in GSH synthesis and cellular redox homeostasis.

### CBX3 binds directly to the promoter of *Slc7a11*, *Slc3a2*, *Ptgs2* and *Cth* to maintains their transcription

The previous bio-informatic analysis has identified *Slc7a11*, *Slc3a2*, *Ptgs2* and *Cth* as potential CBX3 target genes, distinct CBX3 binding peak is discovered within their respective promoter region (**Figure S3)**. To validate this result and determine how the absence of *CBX3* affects their transcription, two complementary chromatin immunoprecipitation (ChIP) assays were performed *in vitro* using HT29 WT and *CBX3*KO cells. ChIP using an anti-CBX3 antibody was carried out to confirm specific CBX3 binding at the promoters of these genes, whereas ChIP with an anti–RNA polymerase II (Pol II) antibody was used to assess the impact of CBX3 deletion on their transcriptional activity. CBX3 and Pol II chromatin association was detected by PCR using specific primers to the identified binding site for each gene. In WT cells, an enrichment of CBX3 and the transcription marker Pol II were identified in the promoter region of these genes. However, in *CBX3* KO cells, the binding of CBX3 is disappeared and the binding of the transcription marker Pol II is decreased on these sites (**Figure 6A-6H**).

**Figure 6:**
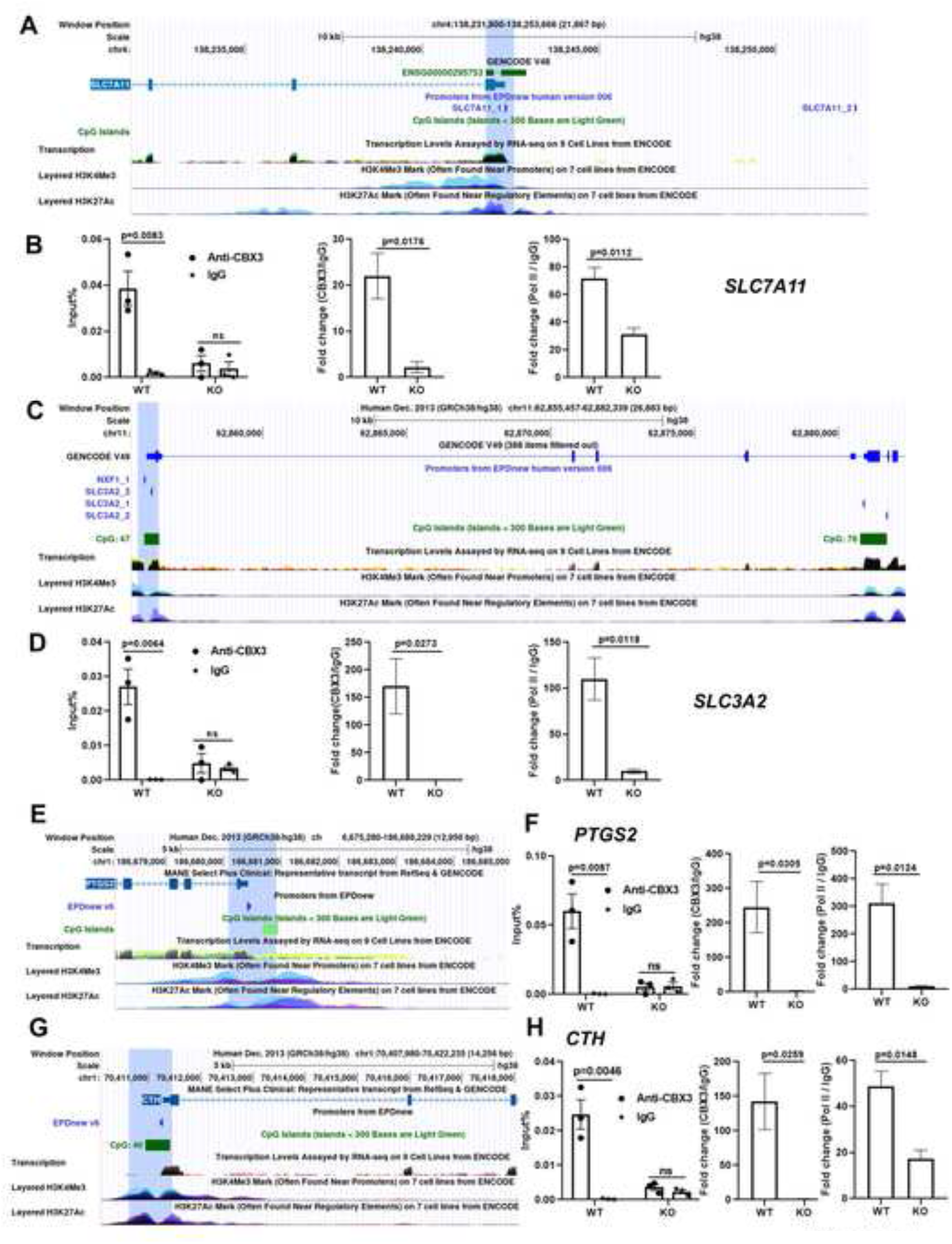
**A, C, E, G**) UCSC Genome Browser centered on SLC7A11 gene (**A**), SLC3A2 gene (**C**), PTGS2 gene (**E**) and CTH gene (**G**): The blue labeling correspond to the different CBX3 binding site of each gene based on the publicly available CBX3 ChIP-seq data performed in HCT116^23^. These binding sites are all located at the gene promoter region enriched by CPG island, H3K4Me3 and H3K27Ac. **B, D, F, H**) **Left and middle:** ChIP analysis was carried out in HT29 WT and *CBX3*KO cells with an anti-CBX3 antibody or IgG to check the CBX3 binding in the promoter of *SLC7A11*(**B**), *SLC3A2*(**D**), *PTGS2*(**F**) and *CTH* (**H**). CBX3 chromatin association was detected by PCR with specific primers to CBX3 binding regions of different gene (bleu labeling in Figure 6A, 6C, 6E or 6G). The left panel represents input% of CBX3 and IgG antibodies and the middle panel represents normalized enrichment (Fold change = Input% of CBX3 antibody / Input% of IgG). Anti-CBX3 ChIP revealed a specific CBX3 binding in the promoter of *SLC7A11*(**B**), *SLC3A2*(**D**), *PTGS2*(**F**) and *CTH* (**H**). However, in CBX3 knockout cells, CBX3 binding at these sites disappeared. Three independent experiments, two-sided Student’s t-test. The error bar represented SEM. **Right)** ChIP analysis with an anti-Pol II antibody or IgG revealed a specific Pol II binding in the promoter of *SLC7A11*, *SLC3A2*, *PTGS2* and *CTH* in HT29 WT cells but this binding is significantly decreased in *CBX3*KO cells. Normalized enrichment (Input% of Pol II antibody compared to Input% of IgG) is presented. Three independent experiments, two-sided Student’s t-test. The error bar represented SEM.

These results indicate that CBX3 directly binds to the promoters of *Slc7a11* (**Figure 6A-6B)**, *Slc3a2* **(Figure 6C-6D)**, *Ptgs2* **(Figure 6E-6F)** and *Cth* **(Figure 6G-6H)**. The CBX3-binding sites on these gene promoters are essential for recruiting Pol II to the same region, thereby maintaining its transcriptional activity in CRC cells.

### *CBX3* deletion-induced downregulation of *SLC7A11* and *SLC3A2* transcription is IFNγ/STAT1 dependent

It was shown that IFNγ signaling could downregulate SLC7A11 via increasing STAT1 binding around the *SLC7A11* transcription-start region^13^. Interestingly, our previous work revealed that *CBX3* deletion could significantly increase transcription and activation of STAT1 under IFNγ stimulation^24^. Therefore, we investigated whether the downregulation of SLC7A11 and SLC3A2 caused by CBX3 deficiency is also STAT1-dependent.

Firstly, western-blot and qPCR analysis (**Figure 7A and 7B**) revealed that *CBX3* deletion significantly decreased SLC7A11/SLC3A2 expression under basal condition. IFNγ treatment decreased SLC7A11 and SLC3A2 expression in both WT and KO HT29 cells, but once more, *CBX3*KO cells still exhibited a lower SLC7A11 and SLC3A2 expression level compared to WT cells. At the same time, higher STAT1 expression and activation level (as indicated by phospho-STAT1 in western-blot analysis) are identified in KO cells compared to WT cells under IFNγ treatment, which is fully consistent with our previous study^24^. Thus, *CBX3* KO-induced IFNγ/STAT1 activation could heighten CBX3 downregulated SLC7A11 and SLC3A2 expression, suggested IFNγ/STAT1 functionally integrated with CBX3 to regulate SLC7A11/SLC3A2 expression.

**Figure 7.**
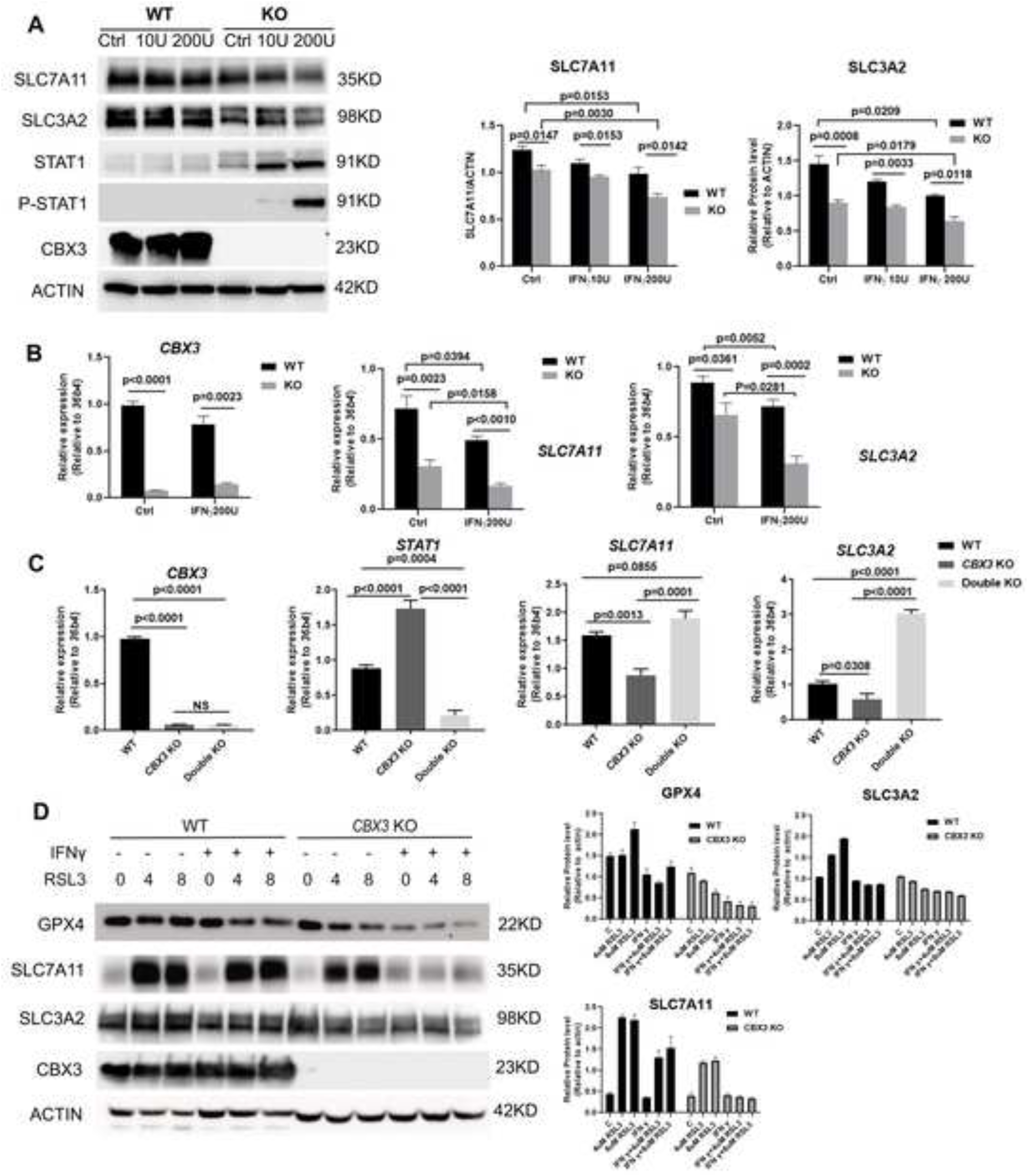
**A**) Western-blot analysis (**left**) and its’ quantification based on the expression level of SLC7A11 and SLC3A2 relative to ACTIN. (**Right**) revealed that KO CBX3 significantly reduced the expression of SLC7A11 and SLC3A2 and IFNγ stimulation could further heighten this inhibition. Two-sided Student’s t-test or two-way ANOVA test, n=3. The error bar represented SEM. B) qPCR analysis revealed that KO CBX3 significantly reduced the transcription of *SLC7A11* and *SLC3A2* and IFNγ stimulation could further heighten this inhibition. **C)** qPCR analysis revealed that single KO of CBX3 (*CBX3*KO) reduced the transcription of *SLC7A11* and *SLC3A2* but double KO *STAT1* and *CBX3* (Double KO) rescued their transcription. **D) Left)** Represented Western blot analysis with WT and CBX3KO HT29 cells showed GPX4, SLC7A11 and SLC3A2 expression under IFNγ, RSL3 or RSL3 plus IFNγ treatment with the same protocol described in Figure2B. IFNγ (200U/ml) and RSL3 was 4 or 8 µM. **Right)** The quantification of the western blot presented in Figure 2C (n=3, the error bar represented SEM) is realized by Image J, based the expression level of GPX4, SLC7A11 and SLC3A2 relative to ACTIN.

We further knocked out *STAT1* gene in *CBX3* KO HT29 cells to check the role of STAT1 in CBX3 KO induced SLC7A11/SLC3A2 downregulation. We successfully performed qPCR analysis in double KO cells 96h after the transfection of a lentivirus contain Crisper targeting STAT1 besides WT and *CBX3*KO cells. As shown in **Figure 7C**, *CBX3* and *STAT1* expression are significantly reduced in double KO cell compared to WT cells. Moreover, *CBX3* single knockout cells significantly reduced the transcriptional levels of *SLC7A11* and *SLC3A2*, but the double knockout of *STAT1* in *CBX3* KO cells restored the transcriptional levels of *SLC7A11* and *SLC3A2*. Therefore, *CBX3* KO-induced downregulation of SLC7A11/SLC3A2 expression was STAT1-dependent.

### *CBX3* deletion impaires the upregulation of SLC7A11/SLC3A2 under RSL3 treatment

We next evaluated how CBX3 deletion affected the expression of GPX4, SLC7A11 and SLC3A2 in CRC cells under RSL3 and IFNγ treatment. (**Figure 7D)**. As revealed by Western blot analysis, compared to WT cells which showed an evident increase of SLC7A11 under RSL3 treatment (alone or with IFNγ), *CBX3*KO impaired RSL3-mediated up-regulation of SLC7A11 and IFNγ plus RSL3 further reinforced this effect. Moreover, SLC3A2 and GPX4 expression is also more evidently reduced in HT29KO cells compared to WT cells under RSL3 treatment and further lessened under IFNγ and RSL3 co-treatment.

The System Xc⁻ (SLC7A11 and SCL3A2) plays a crucial role to maintain redox hemostasis, conferring resistance to ROS stress and ferroptosis. *CBX3* KO impaired SLC7A11, SLC3A2 and GPX4 expression in response to RSL3 plus IFNγ treatment, suggested an increased ferroptosis susceptibility in *CBX3* KO CRC cells.

### Human *CBX3*KO CRC cells are more susceptible to ferroptosis induction, especially combine ferroptosis inducer with IFNγ stimulation

We then evaluated whether knockout *CBX3* also makes human CRC cells more susceptible to two widely used ferroptosis inducers (RSL3 and Erastin), with or without IFNγ stimulation. RSL3 directly binds and inactivates GPX4 and Erastin directly inhibits the system Xc⁻ transporter. Cell viability assay after ferroptosis induction was carried out with CRISPR/Cas9 *CBX3*KO and WT CRC cell lines, SW480 and HT29 (**Figure 8A-8B**).

**Figure 8.**
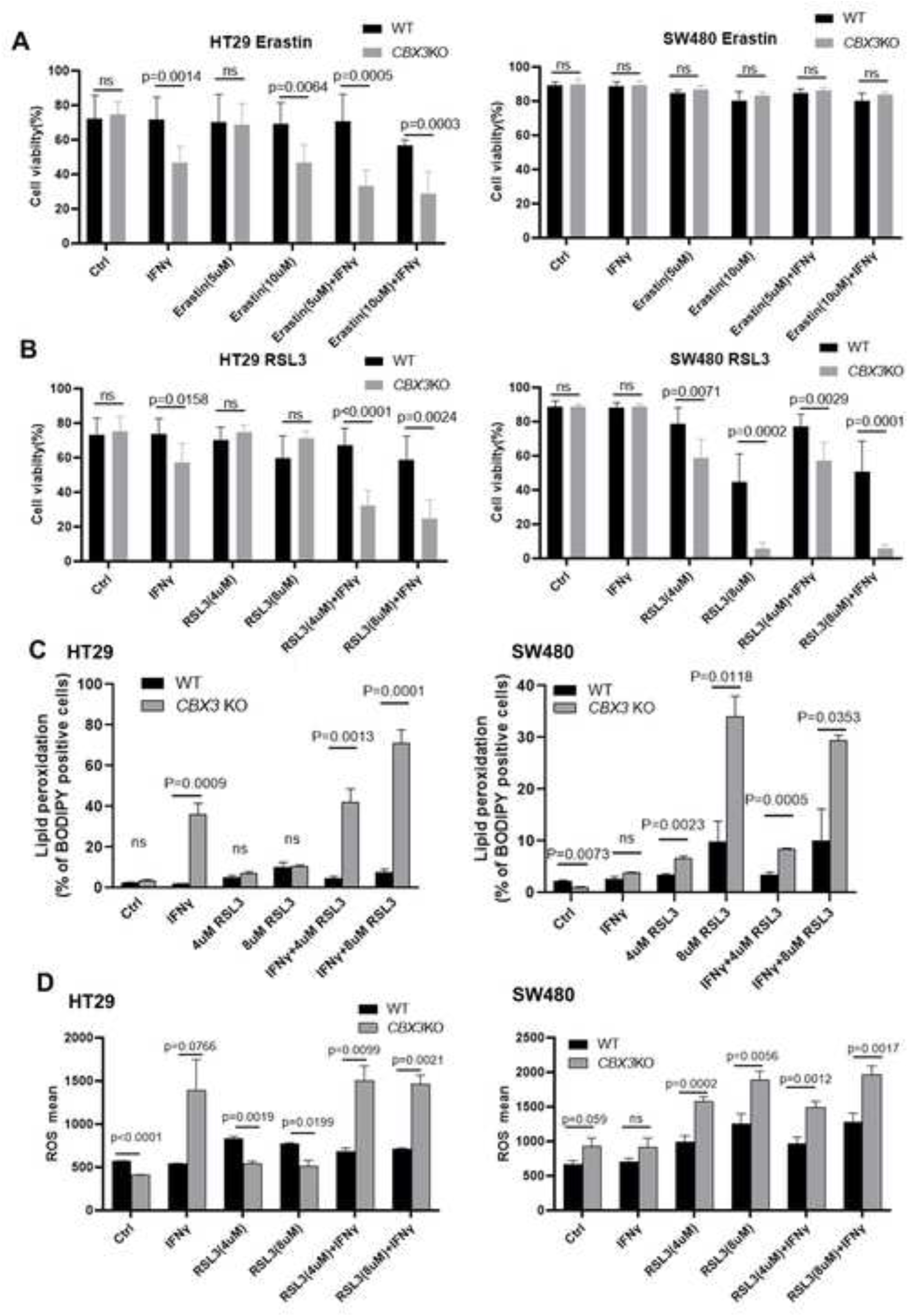
**A**) Left: CBX3KO HT29 cells are more susceptible to ferroptosis inducer Erastin and IFNγ reinforces this effect. **Right:** However, SW480 cells, either *CBX3*KO or WT, exhibited similar resistance to Erastin. The cells were treated with or without human IFNγ(200U/ml), 5 µM or 10 µM of Erastin were added after 24 hours IFNγ treatment and the cells are then cultured for an additional 48 hours, respectively, before analysis. The percent of live cells with different treatments were analyzed by flow cytometry with DAPI labeling indicating dead cells. 6-7 times independent experiments, multiple t test, the error bar represented SEM. **B) Left:** RSL3 plus IFNγ induced more cell death in *CBX3*KO HT29 in a dose dependent manner. **Right:** RSL3 alone is enough to induced more cell death in SW480 cells, but IFNγ cannot further strengthen this effect. After 24 hours with human IFNγ (200U/ml) treatment, 4 or 8 µM of RSL3 were added respectively for an additional 24 hours before analysis. 5- or 6- times independent experiments, two-sided Student’s t test, the error bar represented SEM. **C)** The lipid ROS levels of HT29 and SW480 (WT and *CBX3*KO) cells under IFNγ, RSL3 or RSL3 plus IFNγ treatment with the same protocol described in Figure 8B. Lipid ROS levels were measured with Bodipy 581/591 C11 by flow cytometry under different treatments. 3-8 independent experiments, unpaired two-sided t test. The error bar represented SEM. **D)** The ROS level of HT29 and SW480 (WT and *CBX3*KO) cells under IFNγ, RSL3 or RSL3 plus IFNγ treatment with the same protocol described in Figure 8B. ROS levels of WT and CBX3KO cells were measured with CellROX™ Deep Red probe or CM-H2DCF-DA by flow cytometry. 3-8 independent experiments, unpaired two-sided t test. The error bar represented SEM.

Deletion *CBX3* in HT29 cells alone is sufficient to increase ferroptosis susceptibility to Erastin: the high dose Erastin (10 µM) treatment significantly resulted in higher cell death rates in *CBX3*KO cells compared to WT cells (**Figure 8A, left**). Moreover, IFNγ co-treatment further increased *CBX3*KO CRC susceptibility to Erastin compared to WT cells, consistent with the findings for RSL3: even low dose of RSL3 (4 µM) or Erastin (5 µM) with IFNγ is enough to overcome the ferroptosis resistance of WT cells (**Figure 8B, left**).

Deletion CBX3 in SW480 cells also increased its susceptibility to ferroptosis inducer, but this effect was specifically associated with RSL3 (**Figure 8B, right)** rather than Erastin (**Figure 8A, right)**. Deletion *CBX3* in SW480 cells is sufficient to enhance significantly RSL3 induced cell death in dose dependent manner. However, IFNγ stimulation cannot further enhance this effect.

Briefly, the absence of the CBX3 could generally increases the susceptibility of human CRC cells to ferroptosis. There is a certain degree of cell-specificity probably due to the unique cellular characteristics of SW480 and HT29. As shown by our cytometry analysis, SW480 WT cells exhibits much higher intracellular cystine level compared to HT29 WT cells. Even *CBX3* KO significantly reduced cystine uptake of SW480 cells, the intracellular cystine levels in SW480 KO cells remained higher than those in HT29 WT cells. Consistently, SW480KO cells remain resistant to Erastin, whereas HT29 KO cells do not (**Figure S6 A**). Again, even *CBX3* KO increased ferrous iron loading in both HT29 and SW480 cells, IFNγ only further increased ferrous iron levels in HT29 *CBX3*KO cells but not in SW480 *CBX3*KO cells. Consistently, IFNγ stimulation further enhanced RSL3-induced ferroptosis in HT29 *CBX3*KO cells but not in SW480 *CBX3*KO cells (**Figure S6 B**).

### *CBX3* deletion in CRC cells exacerbates lipid peroxidation and ROS production under RSL3 and IFNγ stimulation

Ferroptosis is characterized by the iron-dependent accumulation of lipid ROS (lipid peroxides), often accompanied by elevated general ROS.

Using BODIPY-C11, a membrane-targeted lipid ROS sensor, we detected changes in lipid ROS accumulation followed different treatments with either RSL3, or IFNγ or combined two of them by flow cytometry analysis. At the same time, we measured also ROS levels in HT29 and SW480 WT and KO cells under different treatments by flow cytometry analysis with CellROX™ Deep Red or CM-H2DCF-DA probe.

The ROS and lipid ROS exhibited similar overall trends. In HT29 cell, combined RSL3 and IFNγ treatment induced a more pronounced increase in lipid ROS and total ROS levels in *CBX3*KO cells compared to WT cells (**Figure 8C).** In SW480 cell, the treatment with RSL3 alone is sufficient to increase significantly lipid ROS and ROS level in *CBX3*KO cells but IFNγ cannot further enhance this effect (**Figure 8D**).

Particularly, only IFNγ but not IFNα could reinforce knockout *CBX3* induced ROS production, suggested that the effect of CBX3 in maintaining redox balance is specifically linked its role to antagonize IFNγ signaling pathway (**Figure S7**).

Overall, our *in vitro* findings indicate that under IFNγ plus RSL3 treatment, *CXB3* deletion in CRC tumor cells exacerbate the ferroptosis hallmarks, namely increased lipid ROS and ROS levels.

## Discussion

Colorectal cancer (CRC) often develops resistance to chemotherapy and other conventional therapies, making alternative cell-death pathways critical. Ferroptosis exploits the redox vulnerabilities of tumor cells, providing a strategy to overcome therapy resistance. The system Xc⁻, a cystine/glutamate antiporter composed of the transporters SLC7A11 and SLC3A2, imports extracellular cystine in exchange for intracellular glutamate. This system supplies cystine for GSH synthesis, maintaining redox homeostasis and protecting cells from lipid peroxidation and ferroptosis.

In this work, we identified that an epigenetic regulator CBX3 is directly implicated in maintain SLC7A11 and SLC3A2 transcription and ferroptosis resistance of CRC (**Figure 9)**. In fact, the presence of *CBX3* is necessary for the binding of POL II to the promoter of *SLC7A11* or *SLC3A2* and maintain their transcription. Deletion of CBX3 downregulated the expression of *SLC7A11*/*SLC3A2.* IFNγ, a cytokine known to promote ferroptosis by downregulating SLC7A11 via STAT1^13^, further heightened these downregulations. Our previous work revealed that CBX3 deficiency enhances the sensitivity of CRC cells to IFNγ via upregulated STAT1 expression^24^. In this work, we further confirmed that the downregulation of SLC7A11 and SLC3A2 induced by *CBX3* KO is also dependent on IFNγ/STAT1 pathway.

**Figure 9.**
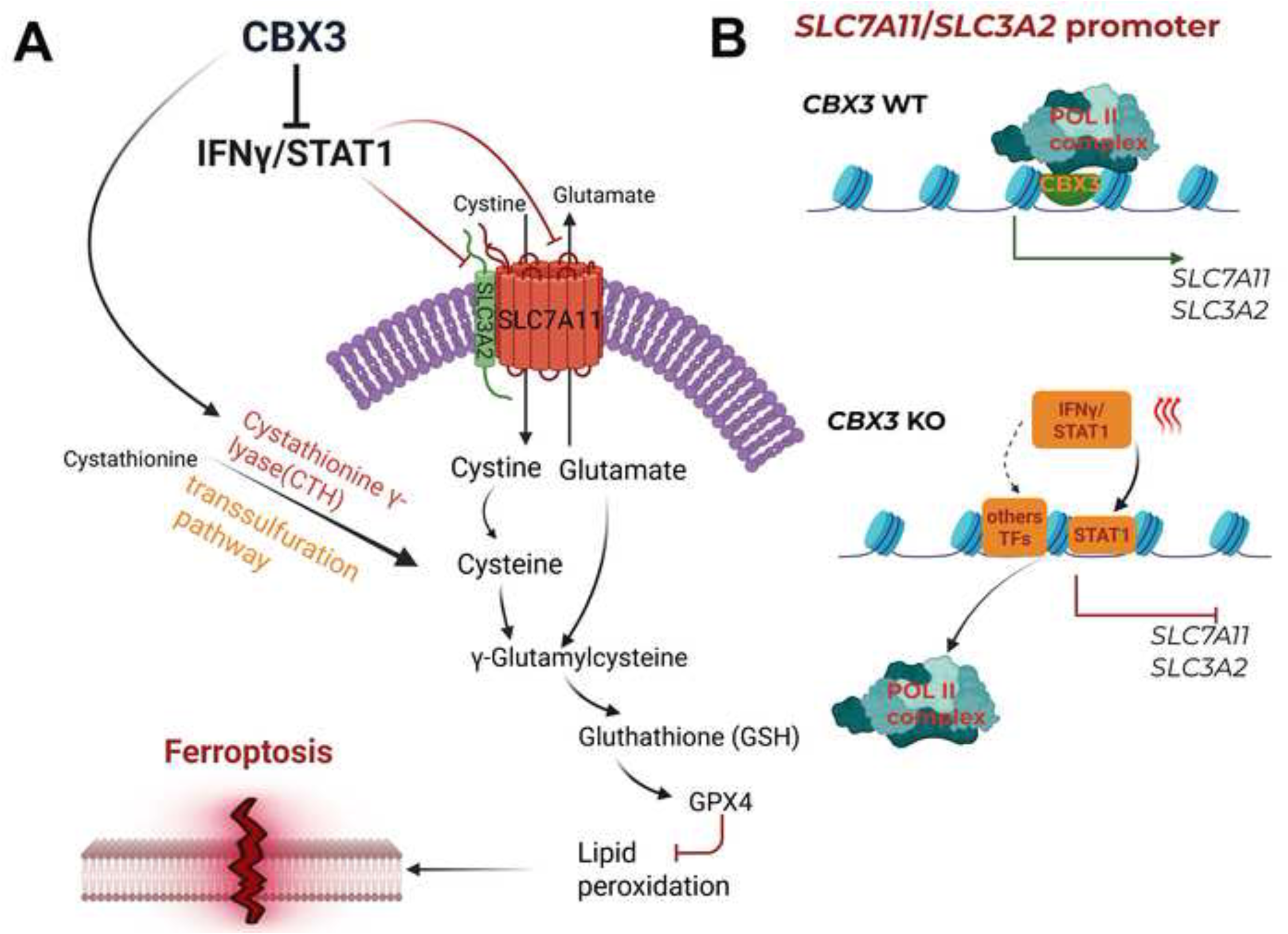
**A**) by Targeting IFNγ/SLC7A11/SLC3A2 Axis, CBX3 is implicated in *SLC7A11* and *SLC3A2* transcription maintenance and ferroptosis resistance of CRC. **B)** the presence of *CBX3* is necessary for the binding of POL II to the promoter of *SLC7A11* or *SLC3A2* and maintain their transcription. Deletion of CBX3 reduced POL II binging to the promotor of *SLC7A11* or *SLC3A2* and downregulated the expression of *SLC7A11*/*SLC3A2.* CBX3 deficiency enhances the sensitivity of CRC cells to IFNγ via upregulated STAT1 expression. The upregulated STAT1 in turn participates the downregulation of SLC7A11 and SLC3A2 induced by *CBX3* KO.

It is noteworthy that although CBX3 deficiency, either with or without IFNγ stimulation, broadly enhances the sensitivity of multiple CRC cells to ferroptosis inducers, this sensitivity is not identical in every respect across all the cells or all the ferroptosis inducers. *CBX3* knockout alone was sufficient to enhance HT29 cells’ sensitivity to Erastin but not to RSL3. In contrast, for SW480 cells, CBX3 loss selectively enhanced sensitivity to RSL3 but not Erastin. Under IFNγ treatment, *CBX3* KO increases HT29 cells’ sensitivity to both RSL3 and Erastin, Whereas IFNγ failed to further sensitize SW480 *CBX3*KO cells to either inducer. *In vivo*, *Cbx3* deletion alone was insufficient to sensitize MC38 tumors to RSL3, only combined use of IFNγ with RSL3 elicited this effect. These differences may originate from the distinct origins of CRC cells, their mutational states, and varying genetic/epigenetic characteristics etc. ^31^ ^32^ ^33^ ^34^. For example, KO *CBX3* makes SW480 cells more sensitivity to RSL3 but not for Erastin. This observation may be related to the KRAS-mutant status of SW480 cells, as KRAS mutations have been reported to promote reliance on GPX4-mediated lipid peroxide detoxification, potentially increasing vulnerability to GPX4 inhibitors such as RSL3 ^35^ ^36^. This increased vulnerability combined with *CBX3* KO finally overcame the resistance of SW480 to RSL3. At the same time, cystine uptake is much higher SW480 cells than HT29 cells. Even *CBX3*KO significantly reduced Erastin uptake in both cell lines but *CBX3*KO SW480 cells still exhibited a higher intracellular cystine level than WT HT29 cells, rendering them resistant to Erastin. Again, as revealed by our previous study, compared to *CBX3*KO HT29 which exhibited an obviously higher phosphorylated STAT1 after IFNγ stimulation, no evident phosphorylated STAT1 was detected in SW480 KO cells despite their remarkable increased STAT1 expression following IFNγ stimulation ^24^. The different STAT1 phosphorylation/activation status could explicate why IFNγ is less effective in SW480 cells compared to HT29 cells and why IFNγ stimulation could further increased the sensitivity of HT29 KO but not SW480 KO cells to RSL3 treatment.

Importantly, CBX3 loss appears globally overcome ferroptosis resistance of different CRC cells regardless their P53 mutant status. In general, CRC with wild-type (WT) p53 is more sensitive to ferroptosis than p53-mutant CRC as WT p53 promotes ferroptosis by repressing SLC7A11^37^. Generally, the MC38 cell line expresses WT P53, whereas the HT29 and SW480 cell line carries a mutant P53 (R273H for HT29; R273H P309S for SW480) ^31^ ^33^. This variance could be a possible reason to explain why WT HT29 cells fail to block the upregulation of SLC7A11 after RSL3 treatment alone or combined with IFNγ, but in WT MC38 tumor, SLC7A11 expression is actually decreased after RSL3+IFNγ treatment. However, knocking out CBX3 in all these cell lines significantly reduced SLC7A11 expression levels, particularly under combined treatment conditions, indicating that CBX3 deficiency enhances the sensitivity of multiple colorectal cancer cell lines to ferroptosis, independent of p53 status.

Beyond SLC7A11 and SLC3A2 which regulate cystine uptake and GSH synthesis, we also identified *Cth* (Cystathionine gamma-lyase) and *Ptgs2* (Prostaglandin G/H synthase 2) are direct targets genes of CBX3. These genes were all downregulated in *Cbx3*KO tumors and further downregulated under combined therapy. CTH catalyzes the last step in the trans-sulfuration pathway from methionine to cysteine. Both SLC7A11/SLC3A2 and CTH are necessary to maintain GSH levels and redox hemostasis, which promotes CRC progression by protecting tumor cells from oxidative stress and ferroptosis ^38^ ^39^ ^40^. Although the exact mechanism of PTGS2 in CRC ferroptosis remains unclear, given that CRC frequently over express PTGS2, which favorited CRC cell proliferation, survival and invasion, and associated with advanced tumor stage, poor differentiation, and worse prognosis, downregulating PTGS2 expression should be a favorable factor for treatment outcomes in colorectal cancer ^41^ ^42^ ^43^ ^44^ ^45^ ^46^. By directly targeting and maintaining their expression, CBX3 may play a crucial role in regulating CRC treatment resistance. Therefore, the expression score of the 11 significantly downregulated genes (Log2FC ≤ −1) in the *Cbx3*KO combo group, including *Slc7a11*, *Ptgs2*, and *Cth*, constitute a prognostic biomarker significantly associated with overall survival in CRC, indicating their importance to evaluate the treatment efficacity.

As the main intracellular antioxidant that protects cells from oxidative stress and helps maintain redox balance, GSH is synthesized in cytosol from Glutamate, Glycine and Cysteine, their uptake/transport in cells depends on several functionally coupled amino acid transporters. Interestingly, *Cbx3*KO significantly reduces the expression of several key amino acid transporters under IFNγ and RSL3 treatment. These amino acid transporters, including SLC3A2, SLC7A5, SLC1A5 and SLC6A9, are functionally coupled with the SLC7A11/GSH axis. The amino acid transporters are essential to sustain CRC high metabolic demands and their survival. The downregulation of several key amino acid transporters implicated in GSH synthesis as a response to oxidative stress, aligns with the suppression of different metabolic pathways critical for lipid and redox homeostasis, reflecting a disorder of cellular metabolism in *Cbx3*KO tumors under combination treatment. Ferroptosis, a metabolism-driven form of regulated cell death, should be highly sensitive to such metabolic perturbations ^8^.

Altogether our findings reveal a critical role for CBX3 in maintaining the functional integrity of IFNγ/ SLC7A11/SLC3A2 axis to protect CRC cells from oxidative stress (**Figure 8D**). By sustaining the expression of key redox and metabolic genes, CBX3 supports ferroptosis resistance, which positions it a promising therapeutic target to enhance the efficacy of ferroptosis-based and redox-targeting therapies in colorectal cancer.

## Abbreviate

CRC: Colorectal carcinoma
CBX3: Chromobox homolog 3
IFNγ: Interferon gamma
ChIP: Chromatin immunoprecipitation
ChIP-seq: Chromatin immunoprecipitation-sequencing
RNAseq: RNA-sequencing
ROS: reactive oxygen species
GSH: glutathione
GPX4: glutathione peroxidase
DEGs: differentially expressed genes
GSEA: Gene set enrichment analysis Human gene will be presented in uppercase italics, e.g., *CBX3;* Mouse genes will be presented in italics with the first letter capitalized, e.g., *Cbx3*, which is also used as a convention for the occasions required indicating both. Human and mouse protein will be presented as uppercase, e.g., CBX3:

## CRediT statements

**Yao XIANG**: Data curation, Formal analysis, Investigation, Methodology, Writing – original draft, Writing – review & editing; **Yuanji FU:** Data curation, Formal analysis, Investigation, Methodology, Writing – original draft, Writing – review & editing; **Christophe Desterke:** Data curation, Formal analysis, Investigation, Methodology, Writing – original draft; **Eric Batsché**: Investigation, Methodology; **Abdelali Jalil:** Investigation; **Rima Elhage:** Investigation, Methodology, Writing – review & editing; **Clémence Duruel:** Investigation; **Jean-Christophe Beche:** Investigation, Resources; **Eric Frapy:** Project administration; **David Skurnik:** Methodology, Writing – review & editing; **Caroline Reisacher:** Methodology, Project administration; **Christine Sedlik:** Methodology, Resources; **Eliane Piaggio:** Methodology, Resources, Writing – review & editing; **Laurence Arbibe:** Methodology, Writing – review & editing; **Ahmed Hamaï:** Data curation, Formal analysis, Investigation, Methodology, Supervision, Writing – original draft, Writing – review & editing; **Yunhua Chang:** Conceptualization, Data curation, Formal analysis, Funding acquisition, Investigation, Methodology, Supervision, Writing – original draft, Writing – review & editing

## Conflict of Interest Statement

The authors declare no potential conflicts of interest

## Supplementary figures

**Figure S1:** HallMARK Geneset GSEA analysis based on WT MC38 tumor RNAseq data revealed that combined RSL3 and IFNγ therapy significantly perturbed oxidative phosphorylation pathways in WT tumors.

**Figure S2:** RNA-seq heatmap generated from RNAseq data exhibited the DEGs with the absolute value of Log2FC≥1 in WT combo group compared to WT Ctrl group.

**Figure S3:** The binding peak of CBX3 is identified in the promotor of *PTGS2*, *CTH*, *SLC7A11* and *SLC3A2* in HCT116 cells by publicly available ChIPseq (Smallwood et al. 2012).

**Figure S4:** The RNAseq data (normalized by EdgeR) revealed the transcription levels of *Slc7a11*, *Ptgs2* and *Cth* in MC38 *Cbx3*KO and WT tumors across the Ctrl and the combo group.

**Figure S5:** Global and individual Schoenfeld tests for covariates included in the Cox Overall survival multi-variable model of TCGA-COAD cohort.

**Figure S6:**

**S6A:** Cytometry analysis using BioTracker Cystine-FITC was performed to analyze Cystine uptake in WT and *CBX3* KO cells. *CBX3* KO significantly reduced Cystine uptake in both SW480 and HT29 cells. But *CBX3* KO SW480 cells still maintain significantly higher intracellular cystine level than WT HT29 cells. (3 independent experiments with SEM., unpaired two-sided t test). The Ratios of fluorescence intensity were calculated using a HT29 control sample as a reference in each independent experiment.

**S6B:** Cytometry analysis using the FerroOrange probe (F374, Dojindo) was performed to analyze ferrous ions in WT and *CBX3* KO cells. Ferrous ion levels significantly increased in two different *CBX3* KO cell types but IFNγ can only further increased Ferrous ion levels in HT29KO cells but not in SW480 KO cells (3 independent experiments with SEM, unpaired two-sided t test). The fluorescence intensity ratios were standardized to HT29 ctrl of every independent experiment.

**Figure S7:** Only IFNγ but not IFNα stimulation could increase ROS production in CBX3KO CRC cells. By flow cytometry analysis with CellROX™ Deep Red probe, ROS production in WT and CBX3KO HT29 was compared under IFNα or IFNγ alone treatment as well as under RSL3+IFNγ or RSL3+IFNα treatment. IFNα alone didn’t increased ROS production in either WT or CBX3KO cells, but IFNγ alone induced significantly a higher ROS production in CBX3KO cells. RSL3+IFNα treatment elevated a similar ROS levels in WT and CBX3KO cells, but RSL3+IFNγ induced significantly a higher ROS production in CBX3KO cells.

## Supplementary material and methods

### Ferrous ion (Fe2+) measurement

FerroOrange probe (F374, Dojindo) were used for the measurement of intracellular iron level. Briefly, HT29 and SW480 cells (30 × 10^3^cells/well) were seeded on a 12 wells plate for two days. Cells were harvested and wished with HBSS three times, then incubated with 1μM FerroOrange in HBSS for 30mins in 37 °C incubator. After labelling, cells were analyzed by LSR Fortessa™ cell analyzer (Becton Dickinson, NJ, USA)

### The Cystine Uptake Assay

HT29 (20000) or SW480 (30000) cells were seeded per well in 12-well plates and cultured for 72 h. Cells were then incubated with 5 μM BioTracker Cystine-FITC (SCT047, Sigma) in cell culture medium for 30 min at 37 °C. Following incubation, cells were washed once with PBS and resuspended in FACS buffer (0,5% BSA, 2mM EDTA in PBS). The mean fluorescence intensity (MFI) was determined by flow cytometry (Sony ID7000) and analyzed using FlowJo 11 software.

## Notes

### Competing Interest Statement

The authors have declared no competing interest.

